# Identifying Ferroptosis Inducers, HDAC, and RTK Inhibitor Sensitivity in Melanoma Subtypes through Unbiased Drug Target Prediction

**DOI:** 10.1101/2024.02.08.579424

**Authors:** Indira Pla, Botond L. Szabolcs, Petra Nikolett Péter, Zsuzsanna Ujfaludi, Yonghyo Kim, Peter Horvatovich, Aniel Sanchez, Krzysztof Pawlowski, Elisabet Wieslander, Jéssica Guedes, Dorottya MP Pál, Anna A. Ascsillán, Lazaro Hiram Betancourt, István Balázs Németh, Jeovanis Gil, Natália Pinto de Almeida, Beáta Szeitz, Leticia Szadai, Viktória Doma, Nicole Woldmar, Áron Bartha, Zoltan Pahi, Tibor Pankotai, Balázs Győrffy, A. Marcell Szasz, Gilberto Domont, Fábio Nogueira, Ho Jeong Kwon, Roger Appelqvist, Sarolta Kárpáti, David Fenyö, Johan Malm, György Marko-Varga, Lajos V. Kemény

## Abstract

The utilization of PD1 and CTLA4 inhibitors has revolutionized the treatment of malignant melanoma (MM). However, resistance to targeted and immune-checkpoint-based therapies still poses a significant problem. Here we mine large scale MM proteogenomic data integrating it with MM cell line dependency screen, and drug sensitivity data to identify druggable targets and forecast treatment efficacy and resistance. Leveraging protein profiles from established MM subtypes and molecular structures of 82 cancer treatment drugs, we identified nine candidate hub proteins, mTOR, FYN, PIK3CB, EGFR, MAPK3, MAP4K1, MAP2K1, SRC and AKT1, across five distinct MM subtypes. These proteins serve as potential drug targets applicable to one or multiple MM subtypes.

By analyzing transcriptomic data from 48 publicly accessible melanoma cell lines sourced from Achilles and CRISPR dependency screens, we forecasted 162 potentially targetable genes. We also identified genetic resistance in 260 genes across at least one melanoma subtype. In addition, we employed publicly available compound sensitivity data (Cancer Therapeutics Response Portal, CTRPv2) on the cell lines to assess the correlation of compound effectiveness within each subtype.

We have identified 20 compounds exhibiting potential drug impact in at least one melanoma subtype. Remarkably, employing this unbiased approach, we have uncovered compounds targeting ferroptosis, that demonstrate a striking 30x fold difference in sensitivity among different subtypes. This implies that the proteogenomic classification of melanoma has the potential to predict sensitivity to ferroptosis compounds. Our results suggest innovative and novel therapeutic strategies by stratifying melanoma samples through proteomic profiling, offering a spectrum of novel therapeutic interventions and prospects for combination therapy.

**Highlights:**

(1) Proteogenomic subtype classification can define the landscape of genetic dependencies in melanoma
(2) Nine proteins from molecular subtypes were identified as potential drug targets for specified MM patients
(3) 20 compounds identified that show potential effectiveness in at least one melanoma subtype
(4) Proteogenomics can predict specific ferroptosis inducers, HDAC, and RTK Inhibitor sensitivity in melanoma subtypes

Graphical abstract

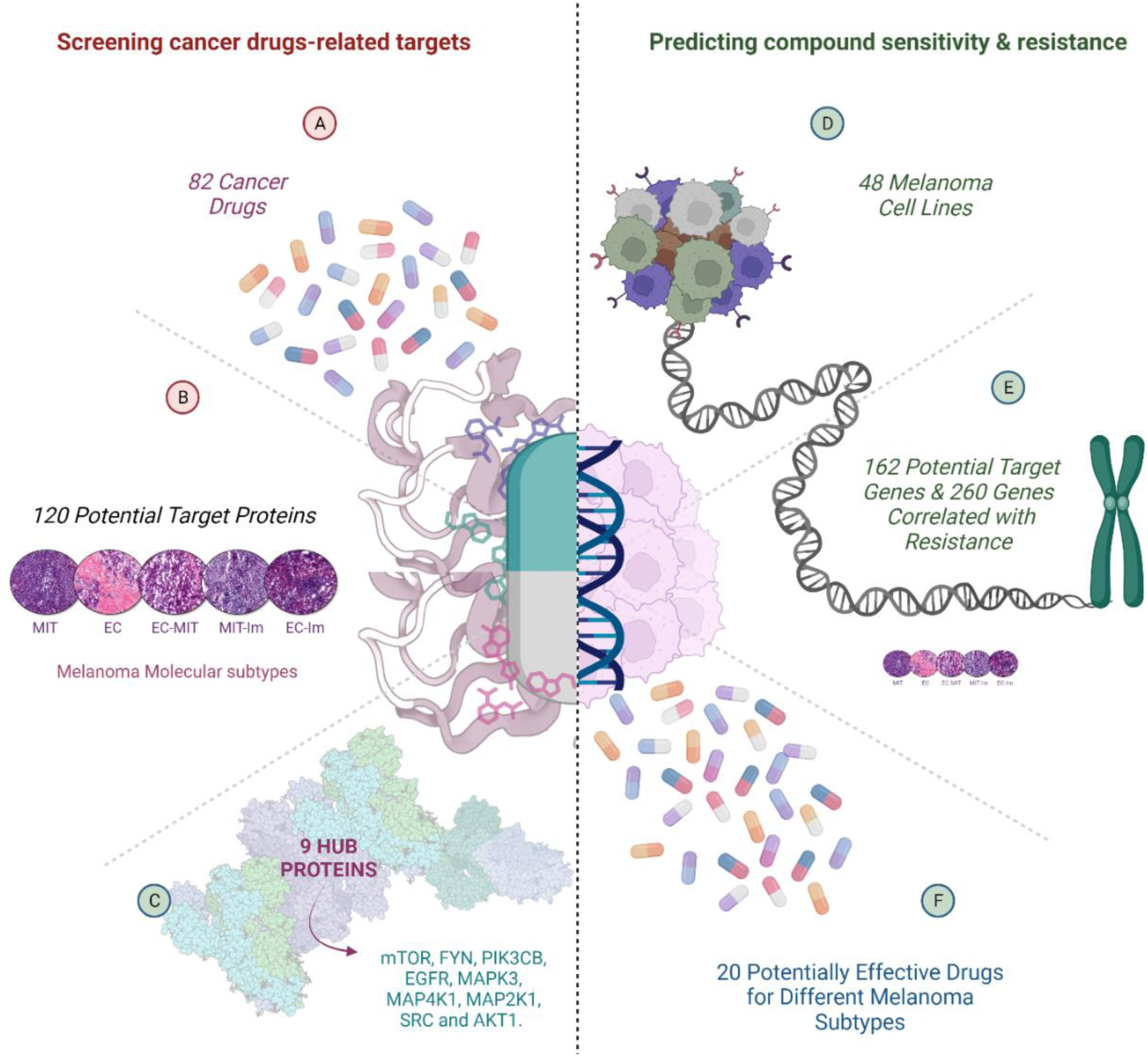

## Introduction

Malignant Melanoma (MM) is the most aggressive skin cancer type and has now become the fifth most common cancer in the US (Cancer Today (iarc.fr)) with more than 300 000 new cases every year (1). The incidence of MM is rising every year, and the estimated number of deaths from MM worldwide is expected to increase by around 57% by 2040 (Cancer Tomorrow (iarc.fr))(2). The appearance of immunotherapies has significantly increased the overall survival of the disease (3). The success of immune-checkpoint inhibitors, such as PD-1 inhibitors and other immunotherapeutic options, has demonstrated the effectiveness of immune reactivation in patients with an advanced stage of the disease. However, MM is also known for being the most heterogeneous cancer type, and despite modern treatments and combination therapies, about half of patients do not achieve long-term improvement. These patients develop resistance to the therapy, and/or get relapse, which induces accelerated progression of the disease. This underlines the necessity of new druggable targets to achieve better results in therapy-resistant patients. It is also essential to discover novel biomarkers to better monitor the therapeutic response (4).

Proteomics plays an important role in discovering new molecules as therapeutic targets since proteins are the main targets of drug discovery (5). Most drugs presently focus on targeting proteins, and within the realm of drug discovery, mass spectrometry (MS)-based proteomics has emerged as a highly valuable analytical method. In addition, this technique has the potential to show the functional status of different subgroups of samples (or individuals) based on high-resolution data, turning it into a cutting-edge technique. In a recent study, our group stratified melanoma metastatic samples into five unique subtypes based on proteogenomic properties, which agrees with this tumor’s well-known heterogeneity (6). Our approach goes beyond classification based on differentiation alone and extends to the analysis of the tumor microenvironment.

Within this study, the five distinct subtypes were: extracellular-immune (EC-Im), mitochondrial-immune (Mit-Im), extracellular (EC), extracellular-mitochondrial (EC-Mit) and mitochondrial (Mit). The first two subtypes (EC-Im and Mit-Im) were associated with high immune activity and a better prognosis, whilst the EC subtype was associated with increased cancer cell proliferation, unfavorable prognosis and short overall survival. Mit subtype exhibited elevated expression of pathways such as “oxidative phosphorylation” and “mitochondrion organization”. Specifically, the EC-Mit subtype displayed notably reduced proliferation compared to the remaining subtypes. This observation raises the possibility that the reduced proliferation rate might be attributed to a dormant stage prompted by the conditions of the host tissue microenvironment. Overall, the Mit and Mit-Im subtypes were related to the occurrence of a more differentiated state, whilst the EC, EC-Im and EC-Mit subtypes were linked to a dedifferentiated state in metastases. Identifying druggable targets specific to subtypes forms the foundation for uncovering novel subtype-related therapies. Furthermore, the anticipation of compound efficacy and assessment of potential resistance holds significant importance to select efficient personalized treatment.

Ligand-based target prediction has exhibited remarkable accuracy in identifying protein targets for biologically active compounds(7,8). Today, bioinformatics tools, such as SwissTargetPrediction (9,10), are accessible for conducting ligand-based target prediction for any small bioactive molecule. On the other hand, the Center for the Science of Therapeutics at the Broad Institute offers a group of tools, such as the Cancer Therapeutics Response Portal (CTRP) to identify and target cancer genetic dependencies (i.e. genes whose products are essential for cancer cell survival) with small molecules (https://portals.broadinstitute.org/ctrp/). This portal is useful for developing insights into small-molecule mechanisms of action and novel therapeutic hypotheses.

In this study, we employed diverse methods to 1) pinpoint drug targets specific to melanoma subtypes, 2) predict the effectiveness and resistance of therapies using transcriptomic profiles from melanoma cell lines and publicly available data from CRISPR edited cancer cell line dependency screens (i.e. data showing survival of cancer cell line after CRISPR gene editing), as well as 3) assess the potency of select pharmacological compounds across different melanoma cell lines.

## Results

This study employed varied approaches and strategies to anticipate subtype-specific targets within melanoma and to gauge sensitivity and resistance to compounds. **Figure 1** shows a summary of the workflow and results that will be described in detail in this section.

**Figure 1.**
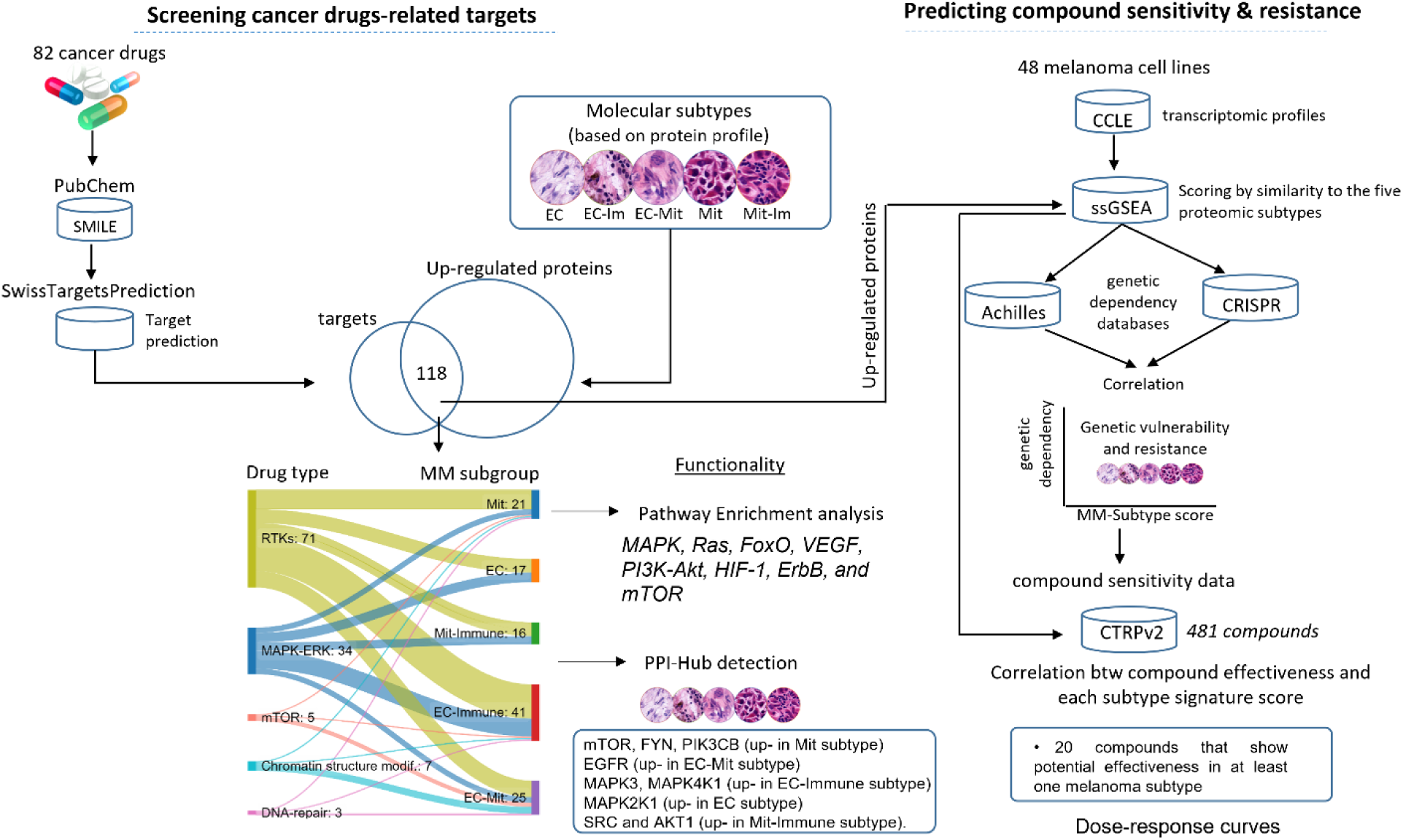
Flowchart of approaches used for screening candidate target proteins, mapping the genetic dependency, and predicting compound sensitivity by pharmacogenomics. The canonical structure of 82 cancer drugs was collected from the PubChem database and uploaded onto the SwissTargetPrediction web service to predict possible protein targets. Putative targets previously reported as melanoma subtype-specific proteins were subjected to pathways enrichment analysis and nine proteins were detected as hubs proteins after performing protein-protein interaction (PPI) analysis using STRING database. Parallelly, the transcriptomic profiles of 48 melanoma cell lines were collected from the Cancer Cell Line Encyclopedia (CCLE), and were scored by similarity to five proteomic subtypes of melanoma using single sample Gene Set Enrichment Analysis (ssGSEA). Then Achilles and CRISPR databases were utilized to detect genetic resistance and vulnerabilities and CTRPv2 database was used for detecting compound sensitivity.

## Protein targets of drugs used in cancer treatments

A comprehensive set of 82 cancer drugs were examined to ascertain potential protein targets (**Table S1**, list of drugs). These were FDA-approved or –investigational drugs known to target receptor tyrosine kinases (RTKs), chromatin structure modifiers, DNA damage, immune checkpoints and the PI3K-Akt, mTOR, Ras and MAPK-ERK signalling pathways, respectively (**Figure 2A**). Through the utilization of the SwissTargetPrediction tool, we identified a total of 118 proteins (as listed in **Table S2**) that exhibited upregulation across the five distinct molecular subtypes of MM patients (as described by Kuras et, al., 2023) (6)and were thus projected as potential targets for the 82 cancer drugs. 63 % of these proteins were protein-modifying enzymes, while the remaining proteins were transmembrane signal receptors (14%), metabolite interconversion enzymes (8%), chromatin/chromatin-binding proteins (5%) and kinases (4%), respectively (**Figure 2B**).

**Figure 2.**
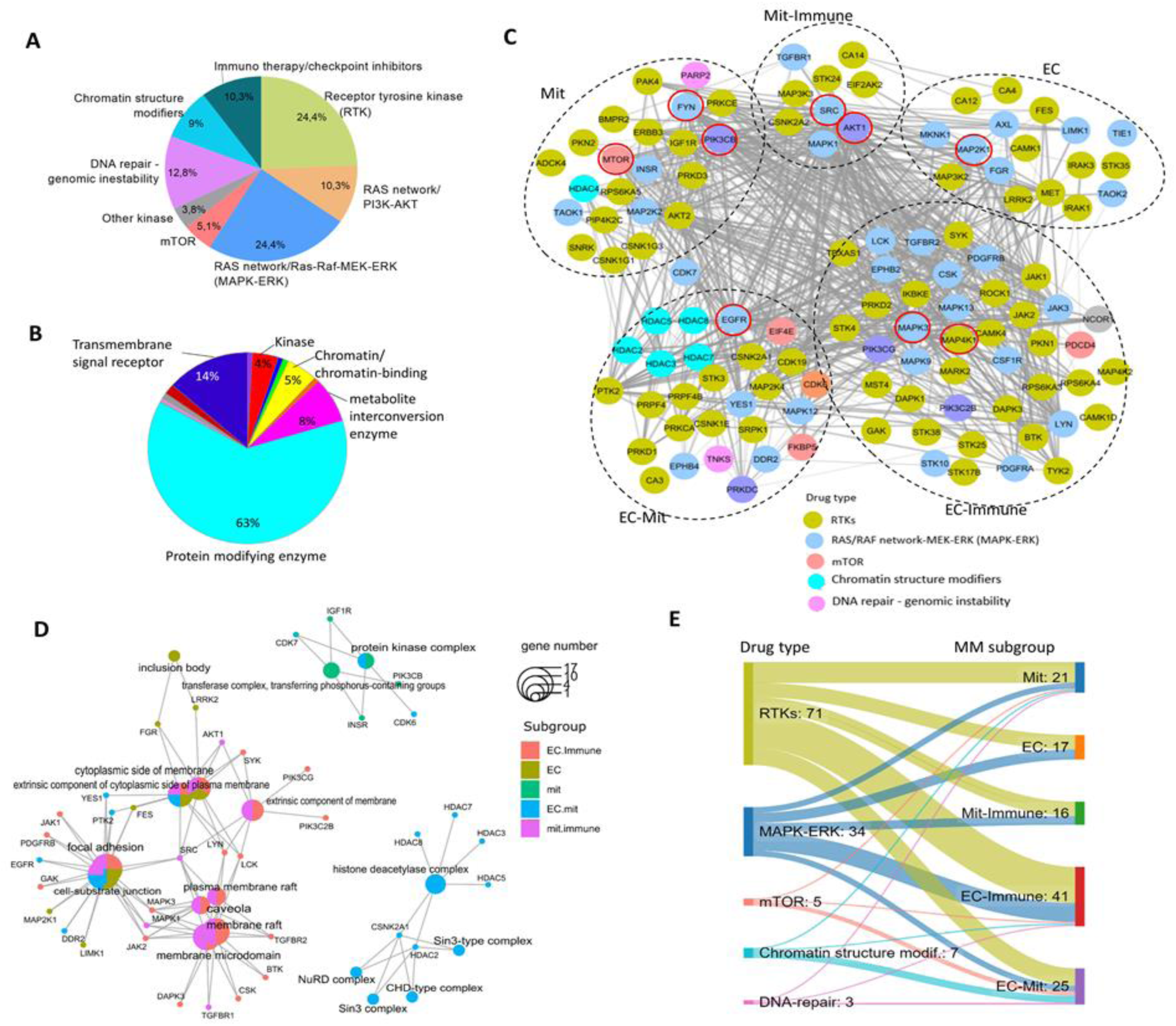
A) Drug types used to identify protein targets in malignus melanoma subtypes. B) Distribution of the ‘protein class’ of the predicted target proteins. C) Functional association network of candidate target proteins upregulated in molecular-defined subtypes of patients with malignant melanoma. Proteins encircled in red color were identified as central network ‘hubs’. D) Cellular component enrichment analysis (FDR < 5%). E) Sankey plot indicating how many proteins upregulated (likely activated) in each MM subtypes were predicted as targets of the different cancer drug types. RTKs: Receptor tyrosine kinases.

Most of the potential targets were discerned through drugs targeting receptor tyrosine kinases (RTKs) (**Figure 2C**). Yet, drugs that alter the chromatin structure and modulators of the mTOR pathway predominantly recognize target proteins activated within a patient subtype previously associated with increased extracellular and mitochondrial protein activity (EC-Mit & Mit) (see **Figure 2C, E**). Upon acquiring the functional association network, (STRING), nine proteins were detected as “hub” proteins (using ‘*cyto*Hubba’ algorithms) (**Figure 2C**). These proteins included mTOR, FYN, PIK3CB, (upregulated and likely active in the Mit subtype), EGFR (upregulated in EC-Mit subtype), MAPK3, MAP4K1 (upregulated in EC-Immune subtype), MAP2K1 (upregulated in EC subtype), SRC and AKT1 (upregulated in Mit-Immune subtype). These hub proteins were estimated as possible targets (*using the <u>SwissTargetPrediction</u> algorithm*) of known cancer drugs (shown in **Table 1**).

**Table 1.** Hub proteins per subtype predicted as possible targets of known drugs.

| Drug/Subtype | EC | EC-Immune | EC-Mit | Mit | Mit-Immune | Total |
| --- | --- | --- | --- | --- | --- | --- |
| Afatinib<br>(TK inhibitor) |  |  | EGFR<br>(0.98) |  | SRC<br>(0.98) | 2 |
| Alpelisib<br>(PI3K inhibitor) |  |  |  | PIK3CB<br>(0.99) |  | 1 |
| Cobimetinib<br>(MEK inhibitor) | MAP2K1<br>($\approx 1.0$) | | | | | 1 |
| Dacomitinib<br>(TK inhibitor) | | | EGFR<br>($\approx 1.0$) | | | 1 |
| Erlotinib<br>(TK inhibitor) |  |  | EGFR<br>(0.98) |  | SRC<br>(0.98) | 2 |
| Gefitinib<br>(TK inhibitor) |  |  | EGFR<br>(*≈1.0*) |  | SRC<br>(*≈1.0*) | 2 |
| Idelalisib<br>(PI3K inhibitor) |  |  |  | PIK3CB<br>(0.75) |  | 1 |
| Imatinib<br>(TK inhibitor) |  |  | EGFR<br>(*≈1.0*) | FYN<br>(*≈1.0*) |  | 2 |
| Ipatasertib<br>(TK inhibitor) |  |  |  |  | AKT1<br>(0.99) | 1 |
| Neratinib<br>(TK inhibitor) | MAP2K1<br>(*≈1.0*) | MAP4K1<br>(*≈1.0*) | EGFR<br>(*≈1.0*) | FYN<br>(*≈1.0*) | SRC<br>(*≈1.0*) | 5 |
| Osimertinib<br>(TK inhibitor) |  |  | EGFR<br>(*≈1.0*) |  |  | 1 |
| Ruxolitinib<br>(TK inhibitor) | MAP2K1<br>(1.0) |  |  |  |  | 1 |
| Selumetinib<br>(MEK inhibitor) | MAP2K1<br>(0.99) |  | EGFR<br>(0.99) |  |  | 2 |
| Sirolimus<br>(Rapamycin)<br>(mTOR inhibitor) |  |  |  | mTOR<br>(*≈1.0*) |  | 1 |
| Sorafenib<br>(TK inhibitor) |  | MAPK3<br>(0.92) | EGFR<br>(0.92) | FYN<br>(0.92) | SRC<br>(0.92) | 4 |
| Sunitinib<br>(TK inhibitor) | MAP2K1<br>(0.99) |  | EGFR<br>(0.99) | FYN<br>(0.99) | SRC<br>(0.99) | 4 |
| TAK632 | MAP2K1<br>($\approx 1.0$) | | | | | 1 |
| Taselisib<br>(PI3K inhibitor) |  |  |  | mTOR;<br>PIK3CB<br>(0.96;<br>0.96) |  | 2 |
| Trametinib<br>(TK inhibitor) | MAP2K1<br>($\approx 1.0$) | | | | | 1 |
| Total | 7 | 2 | 10 | 9 | 7 |  |

Numbers within parentheses indicate the probability (as estimated by SwissTargetPrediction algorithm) of a specific protein to be targetable by the cancer drugs (i.e. small molecule) listed on the left part of the table.
TK: Tyrosine kinase; PI3K: phosphatidylinositol 3-kinase. Color changes according to the scoring with bright red being the best scoring.

Gene ontology (GO) enrichment analysis of cellular component annotations for candidate protein targets revealed that the MM subtypes Mit and EC-Mit displayed enrichment in proteins associated with kinase and histone deacetylase complexes (see **Figure 2D**). In contrast, subtypes of patients associated with high activity of the immune system (EC-Immune & Mit-Immune), were enriched by proteins of the extrinsic component of membrane, plasma membrane raft, and membrane microdomain.

## The functionality of the predicted protein targets

Based on the pathway analysis (shown in **Figure 3**), the candidate targets exhibited enrichment in signaling pathways, including; MAPK, Ras, FoxO, VEGF, PI3K-Akt, HIF-1, ErbB, and mTOR. Pathways associated with T helper cells were enriched within the proteins activated in the subgroup of patients associated with the immune system (EC-Immune & Mit-Immune). Furthermore, the pathway ‘PD-L1 expression and PD-1 checkpoint pathway in cancer’ demonstrated enrichment in proteins activated within the four subgroups associated with elevated activity of mitochondrial and immune-related proteins. Other pathways included ‘EGFR tyrosine kinase inhibitor resistance’, enriched by proteins upregulated in all five subtypes, and ‘MicroRNA in cancer’ enriched in EC_Mit, EC-Immune and Mit subtypes.

**Figure 3.**
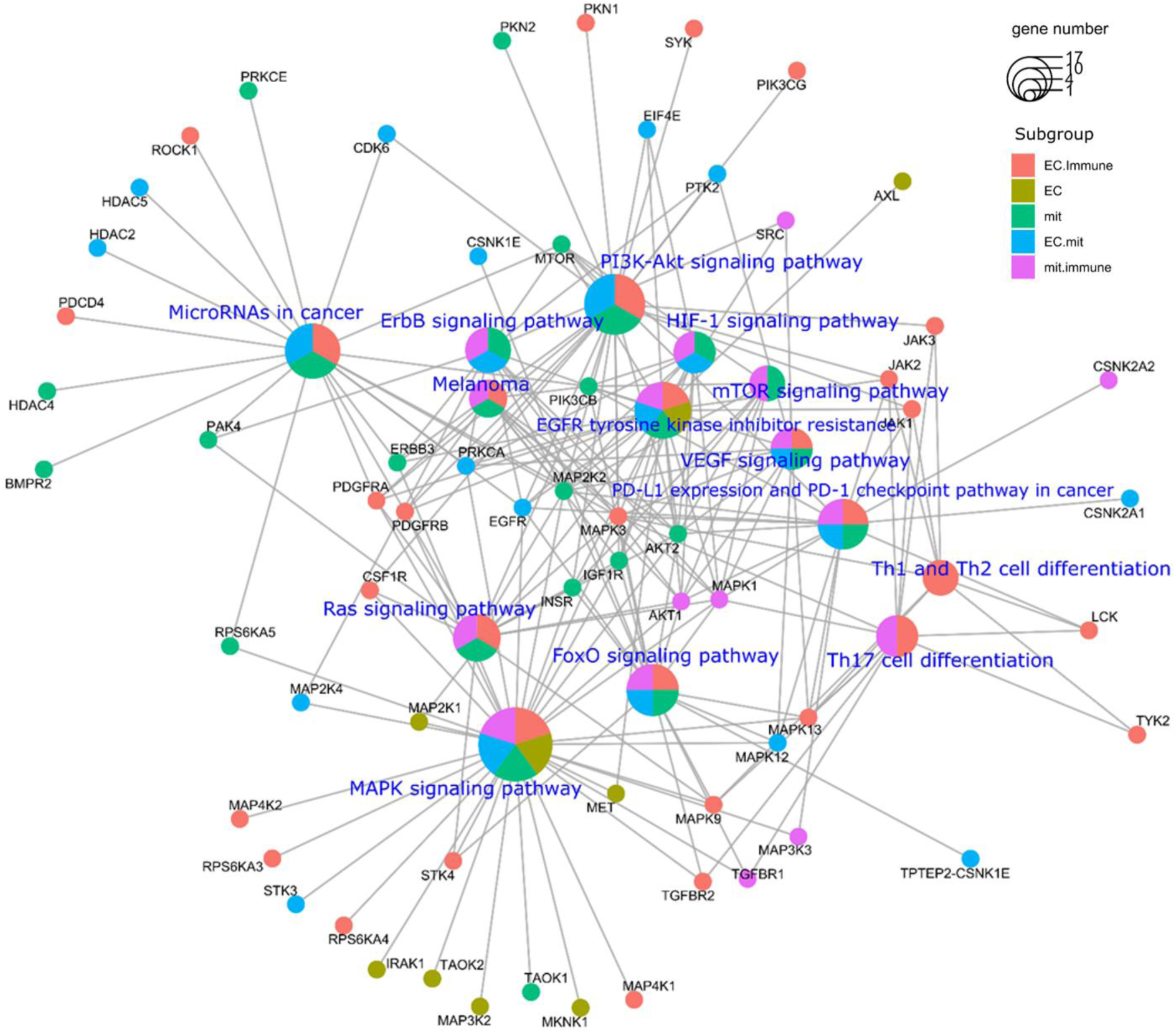
Pathway enrichment analysis based on the KEGG database (FDR < 5%).

## Targetable proteins by subtype

The nine proteins recognized as central hubs within the network of targetable proteins for established cancer treatment drugs were predicted by the ‘SwissTargetPrediction’ algorithm to serve as potential targets for one or several of these drugs (**Table 1**). Table 1 displays the melanoma subtypes showcasing upregulated hub proteins, along with the corresponding drugs through which they were predicted to be targetable. The hub protein MAP2K1 uniquely upregulated in the EC subtype, emerged as a potential target for seven drugs, including Tyrosine kinase (TK) and MEK inhibitors. Proteins MAP4K1 and MAPK3 proteins, upregulated in the EC-Immune subtype, were identified as targets of two TK inhibitor drugs. EGFR, the unique hub protein upregulated in the EC-Mit subtype, was assessed as the target of 10 drugs (mainly TK inhibitors). The three upregulated hub proteins in the Mit subtype (PIK3CB, FYN, mTOR) were predicted as targets for nine drugs. In addition to the nine hub proteins, proteins associated with the histone deacetylase complexes were upregulated and enriched in Mit and EC-Mit subtypes (**Figure 2D**). In particular, histone proteins (HDAC2, HDAC3, HDAC4, HDAC5, HDAC7, HDAC8) exhibited a high likelihood (close to probability of 1) of being targetable by drugs such as Belinostat, Etinostat (excluding HDAC7), Mocetinostat (only HDAC2 and HDAC3), Panobinostat, and Vorinistat. Meanwhile SRC and AKT1 demonstrated upregulation in the Mit-Immune subtype and were identified as targets for 7 drugs. Remarkably, the tyrosine kinase inhibitor Neratinib was found to target at least one hub protein that was upregulated in each of the five subtypes.

## Scoring of melanoma cell lines based on five proteogenomic melanoma subtypes

Subsequently, our objective was to explore the existence of distinct druggable targets within the subtypes, delineated by their expression patterns (this process is illustrated in **Figure 1**). To achieve this, we meticulously analyzed the transcriptomic profiles of 48 melanoma cell lines sourced from CCLE. Each cell line was assigned a score based on its resemblance to the five proteomic subtypes of melanoma, which was accomplished using the single sample Gene Set Enrichment Analysis (ssGSEA) (11,12). Profiles unique to each subtype-specific profiles were characterized by proteins that displayed a significant elevation solely within that particular subtype compared to all other subtypes (as outlined in **Table S2)**.

Following the allocation of subtype-specific similarity scores (ssGSEA scores) to each of the 48 melanoma cell lines, we harnessed publicly accessible genetic dependency databases, namely Achilles and CRISPR. Both datasets are based on CRISPR knockout screens and account for approximately 18,000 genes, which data allows to assess the sensitivity for cell survival of the cell lines to each gene knockout. We established a correlation between genetic dependency across cell lines and each subtypés signature score to ascertain genetic vulnerability (positively correlated) and resistance (negatively correlated) across all five melanoma subtypes (as illustrated in **Figure 4**).

**Figure 4.**
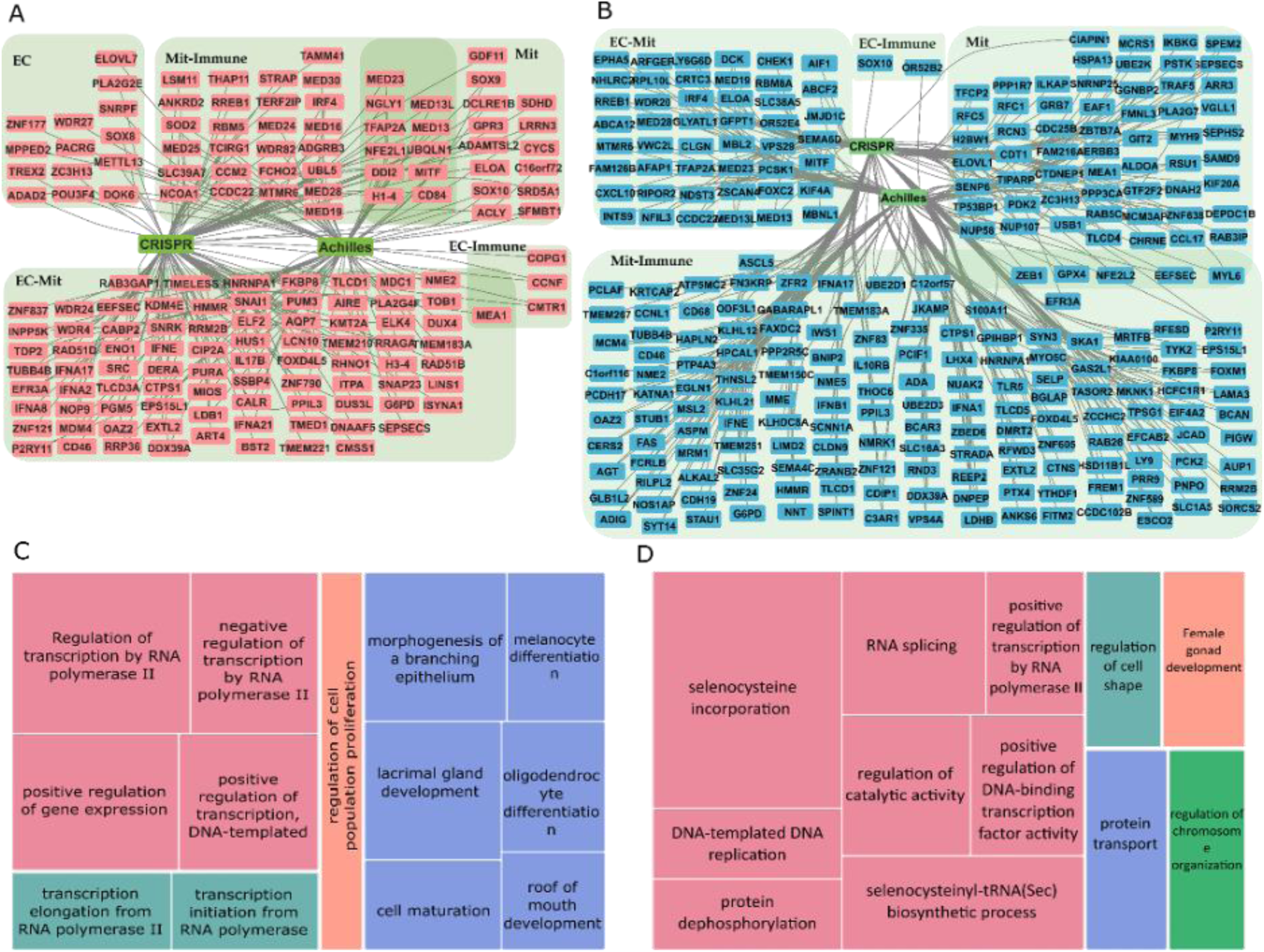
Genetic dependency map of melanoma cell lines with similarity scores to the five melanoma subtypes defined by proteomics. Predicted susceptibility (A, genes indicated with red) and resistance (B, genes indicated with blue) to genetic targeting of individual genes in different melanoma subtypes. Melanoma cell lines from CCLE were scored by similarity with ssGSEA to each melanoma subtype identified previously (6). Across all melanoma cell lines, each subtype signature score was then correlated to the genetic dependency score for each gene obtained from DepMap. Genetic decencies that have demonstrated a significant (FDR < 0.25, p < 0.05) positive (A) or negative (B) Pearson correlation with its corresponding subtype signature score are indicated. Lines connecting individual genes with a CRISPR or Achilles tag indicate the analysis in which the significant association was identified. Pathway analyses (DAVID, followed by REVIGO) of predicted gene dependencies of MIT high and MIT low tumors are displayed in panels (C) and (D), respectively. Note, another interpretation of panels (C) and (D): Pathway analyses of genes that are predicted to show genetic resistance in MIT low (C) and MIT high (D).

Using this approach, we predicted genetic dependencies for each melanoma subtype (depicted in **Figure 4A**, and **Figure S1**) and concurrently identified resistance to genetic knockdown within the various subtypes (depicted in **Figure 4B**, and **Figure S2**). These findings collectively represented potential candidate druggable targets, which includes genetic dependencies and exclude resistant knockout genes. Interestingly, we found that cell lines with Mit high subtype scores displayed sensitivity to the knockout of MITF and SOX10. Intriguingly, cell lines characterized by high Mit-Immune scores also demonstrated susceptibility to the knockout of MITF in addition to IRF4 knockout. These transcription factors – MITF, SOX10 and IRF4 hold pivotal roles in the differentiation and survival of melanocytes and melanoma cells. They are of paramount importance for the survival of well-differentiated tumors (13,14). This may serve as an internal validation for our methodology, given our prior report highlighting the increased differentiation of tumors within the Mit and Mit-Immune subtypes (6). To further support this concept, we extended our investigation by conducting pathway analysis with the genetic dependencies to each subtype (as depicted in **Figure S3**). Our analysis revealed that cell lines characterized by elevated Mit high subtype scores are genetically reliant on genes intricately linked to melanocyte differentiation and cell maturation (illustrated in **Figure 4C**). Conversely, cell lines associated with lower Mit subtype scores demonstrated dependency on genes involved in selenocysteine incorporation (illustrated in **Figure 4D**). These results provide robust support for our methodology and reveal the genetic dependencies of each melanoma subtype.

We have also investigated the melanoma subtypes for potential resistance to genetic knockouts (**Figure 4B**), constructing a genetic resistance map. We observed that cell lines, likely belonging to the EC-Mit and EC-Immune subtypes, displayed resistance to the knockout of the SOX10 MITF and IRF4 genes, aligning with expectations for less differentiated tumors. In accordance with previous observations derived from transcriptomic datasets (13)(Cancer Cell), our investigation unveiled that well-differentiated melanoma cell lines, often attributed to the MIT subtype, exhibited resistance to the knockout of GPX4. This result indicates that specific genetic knockouts may lack effectiveness for distinct melanoma subtypes. However, the data can also be interpreted from another perspective. It implies that cell lines characterized by low Mit subtype scores (indicative of a lower ssGSEA score for MIT signature) exhibit sensitivity to GPX4 knockout. Indeed, the fact that cell lines with Mit and Mit-immune high subtype scores are resistant to genetic deletion of GPX4 suggests that cells with low Mit and Mit-Immune scores (more dedifferentiated) are sensitive to the knockout of GPX4. This aligns with prior observations, as GPX4, a key regulator of ferroptosis, that previously demonstrated efficacy in multi-resistant dedifferentiated melanoma (14).

In summary, using this approach, we projected a cumulative count of 162 genes that hold potential for targeting relevance within at least one melanoma subtype. Concurrently, we identified genetic resistance pertaining to targeting 260 genes across at least one melanoma subtype (refer to **Table 2**, **Table S3, S4**).

**Table 2.**
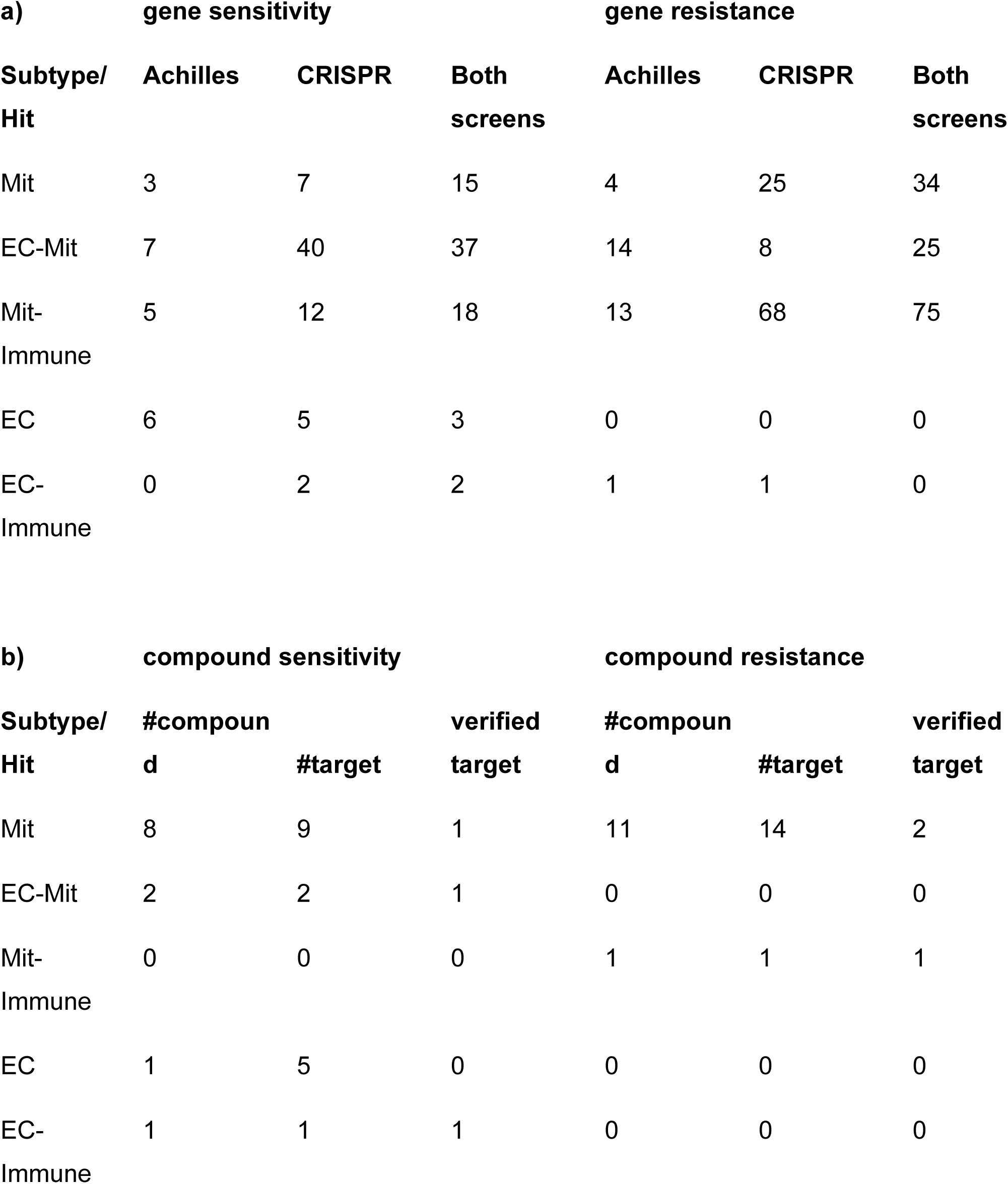
Summary of the number of predicted genetic targets in each melanoma subtype. a) Genetic dependence and b) compound sensitivity of various MM subtypes. Verified targets in panel b indicate the number of target genes of the predicted compounds that show genetic dependency in the dependency datasets as well.

## Compound sensitivity map

Having constructed a dependency map for the melanoma subtypes, our subsequent focus shifted toward evaluating the efficacy of pharmacologic compounds across various melanoma subtypes. To identify pharmacologic compounds, which efficiently kill cells within specific subsets of melanoma subtypes, we utilized a publicly accessible compound sensitivity database (CTRPv2). This database scrutinized the impact of 481 compounds on CCLE melanoma cell lines. We conducted a correlation between potential compound efficacy, as determined by AUC acknowledged for their amalgamation of both efficacy and potency and suggested for superior performance than IC50 values in this dataset (15,16) and each subtype’s signature score using all cell line where the compound was measured (as shown in **Figure 5**). This alignment enabled us to pair compounds with specific melanoma subtypes. We identified 20 compounds exhibiting prospective efficacy (determined by Pearson correlation, FDR < 0.25, p < 0.05) within at least one melanoma subtype (refer to **Table 2, Table S5**). Furthermore, we identified compounds targeting proteins emanating from genes previously identified in the genetic dependency and resistance maps as detailed above. Our analysis highlighted that RSL3 (1S,3R-RSL-3), recognized as a GPX4 inhibitor driving ferroptotic cell death (17), could be effective against cell lines exhibiting elevated scores within the EC-Mit and EC-Immune subtypes (considered as more differentiated subtypes (6). However, its effectiveness appeared to be limited in the other less differentiated melanoma subtypes. We also analyzed the magnitude of the effect between various subtypes on ferroptosis sensitivity. Our findings indicated that contingent on the specific compound, a sensitivity discrepancy ranging from 10 to 50 times was evident between the more differentiated Mit and Mit-immune subtypes compared to the less differentiated subtypes (depicted in **Figure 5C, Table S6**). These results may further indicate the potential utility of ferroptotic compounds in dedifferentiated (EC-Immune, EC-Mit) subtypes. These findings align with our pharmacogenomic-genetic dependency approach detailed earlier and are consistent with previous data derived from transcriptomic datasets and functional experiments (14). Furthermore, our findings suggest the consideration of compounds such as HDAC inhibitors or a combination of carboplatin and paclitaxel. These agents have been previously proposed to exhibit efficacy in melanoma or other cancer types (18,19).

**Figure 5.**
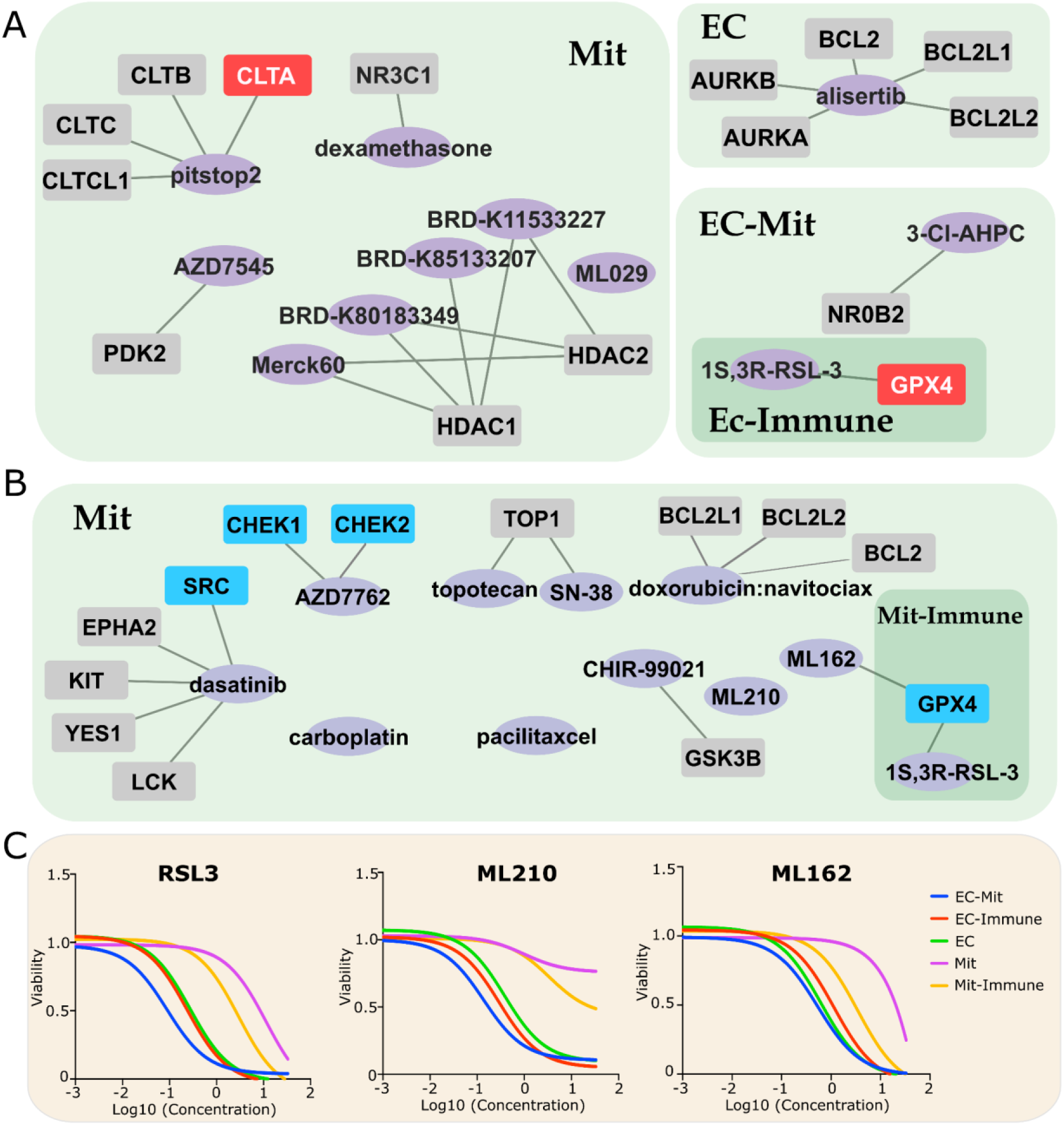
Compound sensitivity map of melanoma subtypes defined by proteogenomics. Predicted sensitivity (A) and resistance (B) to pharmacologic targeting of different melanoma subtypes. Melanoma cell lines sourced from CCLE were assigned scores based on their resemblance to each previous melanoma subtype identified by (6). Each subtype signature score was then correlated to the AUC value of each compound from CTRPv2. Pharmacologic compounds with AUC values that demonstrated significant (Pearson, FDR < 0.25, p < 0.05) positive (A) or negative (B) correlation with its corresponding subtype signature score are indicated. Known targets for each of the compounds that have demonstrated significant correlation with its subtype-specific signature score are indicated with red, blue, or grey for positive, negative or non-significant correlation, respectively. (C). Dose-response curves (raw values obtained from deposited public data) of different ferroptotic compounds demonstrate the magnitude of difference between various subtypes.

To validate our approach, we have performed wet-lab experiments and have observed increased sensitivity for several compounds. Specifically, we decided to validate the potential use of ferroptotic compounds, as indicated by both the genetic dependency map and the compound sensitivity map. As expected, we have seen that cell lines with low MIT signature score (LOXIMVI, A375) demonstrated increased sensitivity to all tested compounds (erastin, RSL3, ML210) than cell lines with higher MIT signature scores (SKMEL28, SKMEL30) (Supplementary **Figure S4**). We also validated the efficacy of Dasatinib in cell lines with low MIT scores (Supplementary **Figure S4**), as Dasatinib has been used in clinical trials and have demonstrated relatively poor responses (20,21) however, in subsets of patients where Dasatinib has been shown to have effectiveness, novel biomarkers may stratify patients by response to das Dasatinib. Our results could potentially stratify patients who may benefit from clinically available drugs that have failed to show overall effectiveness in large populations. Through proteogenomic classification, these therapies could be targeted towards patients who have not responded to modern targeted or immune-based therapies.

## Discussion

Several studies have addressed the categorization of melanoma. Traditionally, melanomas have been classified into subtypes based on the origin of the primary tumor (3). Based on gene expression profiles, Jönsson and colleagues (22) categorized metastatic melanomas into four subgroups: MITF-low/proliferative, high-immune response, MITF-high/pigmentation and normal-like. Moreover, Tsoi et al. identified four metastatic melanoma subtypes at the gene level, correlated with sensitivity to iron-dependent oxidative stress and cell death termed ferroptosis (14). Another avenue of exploration is predicated on driver mutations, while numerous additional investigations are rooted in transcriptomic signatures (22–26).

Applying the proteogenomic classification as described in our recent study (6), we have uncovered that Mit and Mit-Immune subtypes are predicted to be sensitive to the knockout of MITF and SOX10. This aligns with expectations, given the roles of MITF and SOX10 in regulating survival and differentiation. However, SOX10 regulates PD-L1 expression in melanoma and makes the tumor cells less vulnerable to PD-1 inhibitors (25). SOX10 regulates PD-L1 expression through a pathway including IRF4, a pigmentation-associated gene (27,28). According to our results, IRF4 emerges as a significant player in the survival of Mit-Immune cell lines. Our results suggest that directing interventions towards SOX10 signaling within the Mit and Mit-Immune subtypes may be a beneficial strategy due to decreasing melanoma survival. However, it’s important to acknowledge that this strategy might also heighten susceptibility to immune checkpoint blockade *in vivo*, as outlined by Yokoyama et al., 2021. MITF is also regulated by SOX10, and it has been associated with resistance to targeted therapies (27).

GPX4 has a crucial role in the survival of specific melanoma cells (17). Knocking out GPX4 induces the accumulation of lipid peroxides, which leads to ferroptosis. Ferroptosis is an iron-dependent form of cell death (29). Within our study, the compound sensitivity map revealed that cell lines with EC-Mit and EC-Immune high scores are predicted to be sensitive to ferroptosis and ferroptosis-inducing drugs. It is also revealed that the Mit and Mit-Immune high-score cell lines are resistant to ferroptosis. Previous studies have shown similar correlations using transcriptomic datasets and also by functional experiments modifying the differentiation status of melanoma cells (14). Well-differentiated melanomas were proposed to be resistant to ferroptosis-inducing drugs RSL3, ML162, and ML210, unlike undifferentiated melanomas, which were sensitive to them (14). Collectively our results substantiate the potential effectiveness of ferroptosis-inducing drugs in a significant proportion of melanomas.

However, melanoma exhibits substantial heterogeneity leading to varying states of cellular differentiation. A previous study revealed that co-treatment of cancer cells with ferroptosis-inducing drugs and immunotherapy effectively reduces the number of residual persister cells (as demonstrated by (17)). Therefore, combination therapies that include ferroptosis-inducing drugs may be effective in the future, since ferroptosis-inducing drugs can be used to target both dedifferentiated and persistent cells. Hence, future therapeutic strategies that encompass combination therapies featuring ferroptosis-inducing drugs could hold significant promise. These drugs possess the potential to effectively target both dedifferentiated cells and persister cells, offering a comprehensive approach to addressing the complexity of melanoma.

Furthermore, we have successfully identified compounds of interest, including HDAC inhibitors, as well as carboplatin and paclitaxel – all of which have been previously suggested as treatment modalities in melanoma and other cancer contexts. For instance, HDAC inhibitors, like Vorinostat have received FDA approval for treating cutaneous T-cell lymphoma and certain hematologic malignancies (30). A phase II trial demonstrated that Vorinostat showed disease stabilization in a cohort of patients with cutaneous or ocular melanoma(31,32). In parallel, our findings suggest that this combination might hold greater promise within the Mit low subtype of melanoma. It would be interesting to investigate the efficacy of these compounds in Mit low subtypes, as well as the efficacy of HDAC inhibitors in the Mit high subtypes. Furthermore, the fusion of HDAC inhibitors with immune checkpoint inhibitors has been suggested as a combined therapeutic approach to overcome resistance in malignant melanoma cases.

In summary, the recent introduction of proteomic-driven subtype-specific classification for melanoma has demonstrated the diversity of possible approaches available to target this disease. Through the stratification of melanoma samples based on proteomic profiles predicting drug efficacy, our findings offer innovative therapeutic possibilities. However, there are limitations related to our model. Although we used proteogenomic expressions from ex vivo melanoma samples, some of our approaches relied on using cell lines for predicting vulnerabilities which may not be able to recapitulate the full tumor microenvironment. Therefore, it becomes imperative to undertake further wet-lab experiments using short-term melanoma cultures derived from previously proteogenomic-characterized melanoma tumors.

## Material and Methods

### Identification of candidate targetable proteins

Using the SwissTargetPrediction tool (9,10), we predicted the most probable protein targets of drugs currently used in the clinic for malignant melanoma and other cancer types (**Table S1**, list of the drugs). We included FDA-approved or –investigational drugs known to target receptor tyrosine kinases (RTKs), chromatin structure modifiers, DNA damage, immune checkpoints and the PI3K-Akt, mTOR, Ras and MAPK-ERK signaling pathways. The canonical SMILES (simplified molecular input line entry specification) structure of the drugs was collected from the PubChem database(33) and uploaded onto the SwissTargetPrediction web service to predict possible targets. This tool estimates the most probable macromolecular targets of a small molecule assumed as bioactive. The prediction is assessed by searching for similar molecules, in 2D and 3D structures, within a larger collection of compounds known to be experimentally active on an extended set of macromolecular targets (9,10).

### Subtype-specific candidate druggable targets

To establish a connection between the candidate targets and various subtypes of patients with malignant melanoma, we merged the list of predicted targets with the list of proteins previously identified as likely activated (given their upregulation, within the context of five different molecular subtypes of MM patients (n = 142) (6)). The functional association network of the candidate targets was obtained using the STRING online tool (https://string-db.org/), and the ‘hubs’ were estimated using the ‘*cyto*Hubba’ package (34) included in Cystoscape software (35). The topological analysis method selected in ‘*cyto*Hubba’ was Maximal Clique Centrality (MCC). GO and KEGG pathway enrichment analysis was performed using the R package, ‘clusterProfiler’ (36).

### Drug sensitivity/resistance map of melanoma based on proteogenomic profiling

We examined transcriptomic profiles of 48 melanoma cell lines from the cancer cell line encyclopedia (CCLE) (37), and scored each by similarity to the five proteomic subtypes of melanoma using ssGSEA (https://github.com/GSEA-MSigDB/ssGSEA-gpmodule). We used the uniquely overexpressed proteins in each subtype (**Table S2**) to assign subtype-specific signature score to each melanoma cell line (**Table S6**).

### Genetic dependency map

We utilized publicly available data from the genetic dependency databases CRISPR and Achilles, in which the genetic dependency of 18000 genes was studied by CRISPR-mediated knockout (38–40). Using the Morpheus web tool (https://software.broadinstitute.org/morpheus), across all available melanoma cell lines (60 cell lines) each subtype signature score (5 subtype scores for each cell line) was correlated (Pearson correlation) to all genetic dependency scores (20000 genes for each cell line) obtained from DepMap (https://depmap.sanger.ac.uk/), which resulted in 5×20000 correlation pairs between mRNA levels and subtype scores in 60 cell lines). DepMap Public 22Q1 version of CRISPR and Achilles gene dependency datasets were used. Genetic dependencies were considered statistically significant and have demonstrated a significant positive (genetic susceptibility/vulnerability) or negative (genetic resistance) Pearson correlation (FDR < 0.25, p < 0.05) with its corresponding subtype signature score.

### Compound sensitivity map

CTRPv2 drug susceptibility database (41) was used to investigate the relationship between the drug susceptibility and subtype-specific signature score of cell lines. From the CTRPv2 database, we extracted the area under the curve (AUC) value of each compound as a measure of drug response. Each subtype signature score was correlated to the AUC value of each compound from CTRPv2 and pharmacologic compounds with AUC values that demonstrated significant (Pearson, FDR < 0.25, p < 0.05) positive (a) or negative (b) correlation with its corresponding subtype signature score were considered as potential hits. For each compound associated with each subtype, where the protein targets of the compound are identifiable, the corresponding gene is indicated only if its genetic dependency correlated with the subtype signature score (i.e., if the genetic ablation of a particular gene is predicted to be efficient in the subtype where the compound has been predicted to be efficient).

### Pathway analyses of predicted gene targets of each subtype

DAVID bioinformatic web tool (42,43)was used to perform gene set enrichment analysis involving potential target transcripts of the given subtype. REVIGO (44) with SimRel semantic similarity measure and with allowed similarity of 0.5 was used to remove redundant gene sets using the significantly enriched gene sets (p = 0.05) by DAVID.

### Cell lines and culture conditions for validation of predicted drug efficacy

Human melanoma cell lines SKmel-28 (#HTB-72 and A375 (#CRL-1619) were obtained from American Type Culture Collection (ATCC, USA). SKmel-30 cell line (#ACC 151) were purchased from DSMZ-German Collection of Microorganisms and Cell Cultures GmbH. LOX-IMVI (#SCC201) cell lines were purchased from Sigma-Aldrich. The cells were cultured in RPMI1640 with 10% fetal bovine (FBS), and 1% Penicillin/Streptomycin. All cell lines were cultured in a humidified incubator at 37°C with 5% CO_2_. Dasatinib (SML2589) was obtained from Sigma Aldrich, Trametinib, ML210 (#S0788) and RSL3 (#S8155) (GSK1120212) from Selleckchem. Erastin (#HY-15763) was purchased from MedChemExpress (MCE). Melanoma cell lines were seeded in a 96 well plate at a density of 2500 cells/well in 100 ul of complete RPMI1640 and allowed to adhere overnight at 37°C with 5% CO_2_.The next day all cell lines were treated with the compounds at maximal concentration of 32uM (or 3uM for Dasatinib). 1:3 serial dilution was then used for all other doses. After three days of treatment cell viability was measured with CellTiter-Glo Assay (Promega) according to the manufacturer’s instructions and a microplate reader (CLARIOstar Plate Reader) was used to measure viability. Data was analyzed with GraphPad Prism 9 to calculate IC_50_ values, by fitting via nonlinear regression.

## Data Availability Statement

The scripts used for the statistics are available at https://github.com/indirapla/MM500_Duggable_proteome.

## Acknowledgement

This study was supported by grants from the Berta Kamprad Foundation, Lund, Sweden. The generous support by Thermo Scientific, Liconic UK, and Tecan was absolutely key for this study. This work was done under the auspices of a Memorandum of Understanding between the European Cancer Moonshot Center in Lund and the U.S. National Cancer Institute’s International Cancer Proteogenomic Consortium (ICPC). ICPC encourages international cooperation among institutions and nations in proteogenomic cancer research in which proteogenomic datasets are made available to the public. This work was also done in collaboration with the U.S. National Cancer Institute’s Clinical Proteomic Tumor Analysis Consortium (CPTAC). Further support came from the National Cancer Institute (NCI) CPTAC grants U24CA210972 to D.F. I.P. work was funded by the Elisabeth and Alfred Ahlqvist scholarship from the Swedish Pharmaceutical Society. L.V.K. is a recipient of the János Bolyai Research Scholarship of the Hungarian Academy of Sciences and is supported by the Hungarian National Research, Development and Innovation Office (OTKA FK138696) and Semmelweis University STIA-KFI2021 grants. I.B.N. was supported by the Hungarian Academy of Sciences, the grant of OTKA-125509. L.S. was supported by the ÚNKP-21-3-SZTE-102 New National Excellence Program of the Ministry for Innovation and Technology from the source of the National Research, Development and Innovation Fund (University of Szeged, Szeged, Hungary). The project has received funding from the EU’s Horizon 2020 research and innovation program under grant agreement No. 739593.

## Author contribution

Study Conception & Design: I.P., P.H., G.M., L.V.K.

Performed Experiment or Data Collection: I.P., B.L.S., P.N.P., Z.Ú., Y.K., P.H., A.S., D.M.P., A.A.A., L.V.K.

Computation & Statistical Analysis: I.P., B.L.S., P.N.P., A.S.

Data Interpretation & Biological Analysis: I.P., B.L.S., P.N.P., Z.Ú., P.H., J.G., G.D., G.M., L.V.K.

Writing – Original Drafts: I.P., B.L.S., P.N.P., G.M., L.V.K.

Writing – Review & Editing: I.P., B.L.S., P.N.P., Z.Ú., Y.K., P.H., A.S., K.P., E.W., J.G.,

P.M.D., A.A.A., L.H.B., I.B.N., J. Gil, N.A., B.S., L.S., V.D., N.W., Á.B., Z.P., T.P., B.G.,

A.M.S., H.J.W., G.D., F.N., R.A., S.K., D.F., J.M., G.M., L.V.K. Supervision: P.H., K.P., D.F., G.M., L.V.K.

Administration: R.A.

*Suppl. Files 1-6. Pathway analysis of the Biological Processes Gene Set (by DAVID) of genes with genetic dependencies in MIT high, MIT low, MITIMMUNE high, MITIMMUNE low, ECMIT high, ECMIT low*

## Supplementary Figures

**Figure S1.**
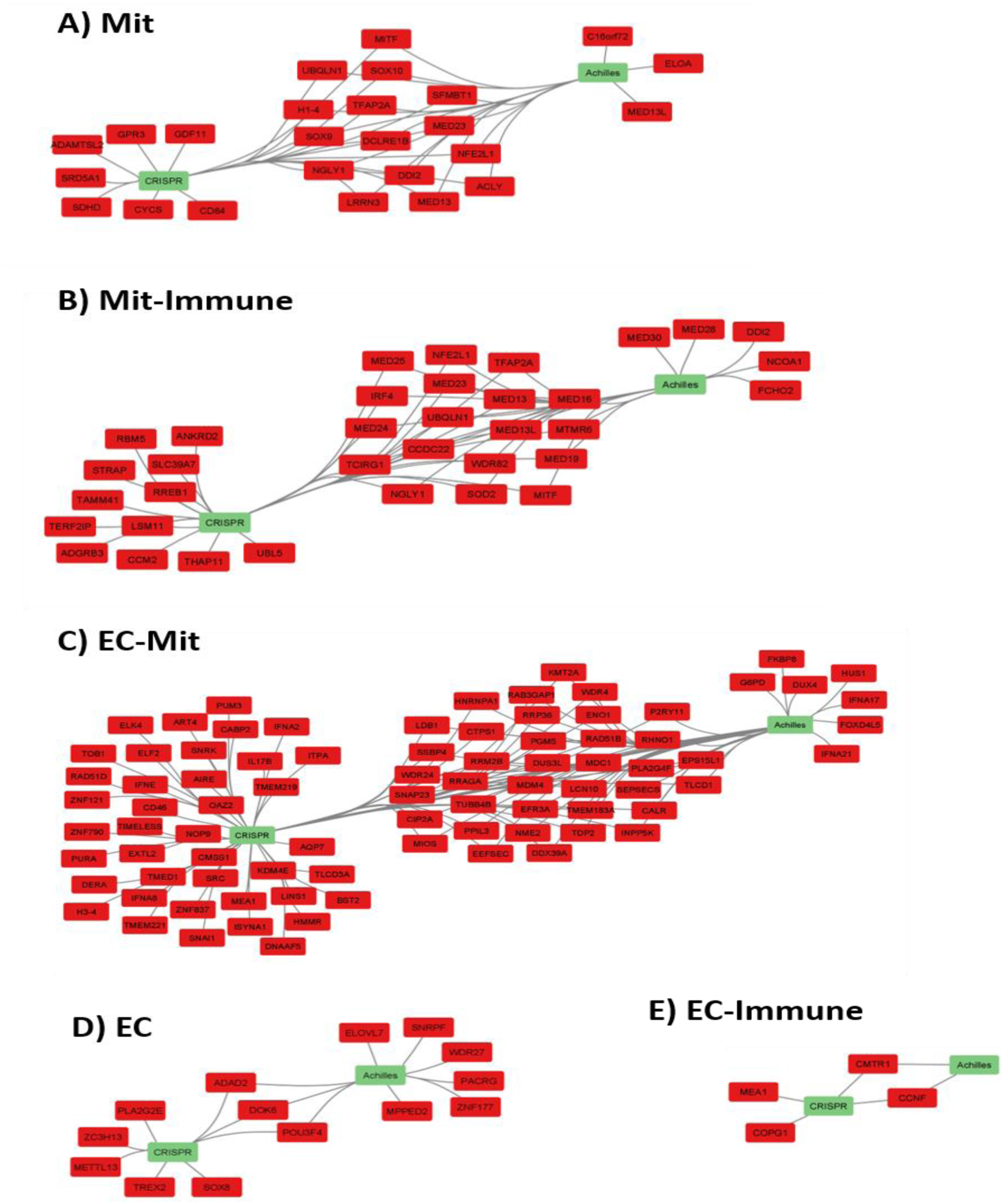
Genetic susceptibility map of melanoma subtypes defined by proteogenomics. Predicted susceptibility to genetic targeting of individual genes in different melanoma subtypes. Methods described in details in Fig. 1. Legend. Lines connecting individual genes with a CRISPR or Achilles tag indicate the analysis in which the significant association was identified

**Figure S2.**
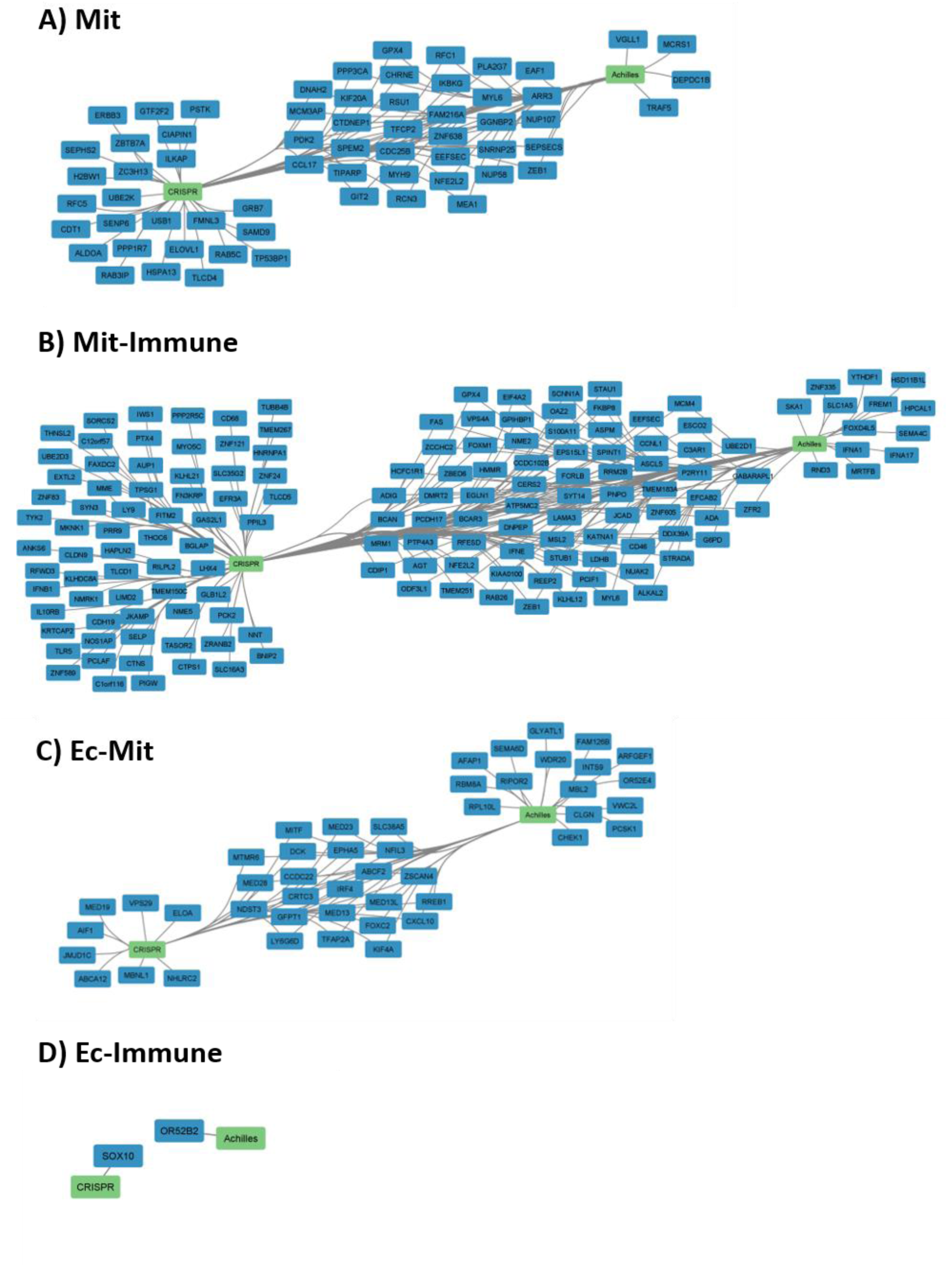
Genetic resistance map of melanoma subtypes defined by proteogenomics. Predicted resistance to genetic targeting of individual genes in different melanoma subtypes. Methods described in details in Fig. 1. Legend. Lines connecting individual genes with a CRISPR or Achilles tag indicate the analysis in which the significant association was identified

**Figure S3.**
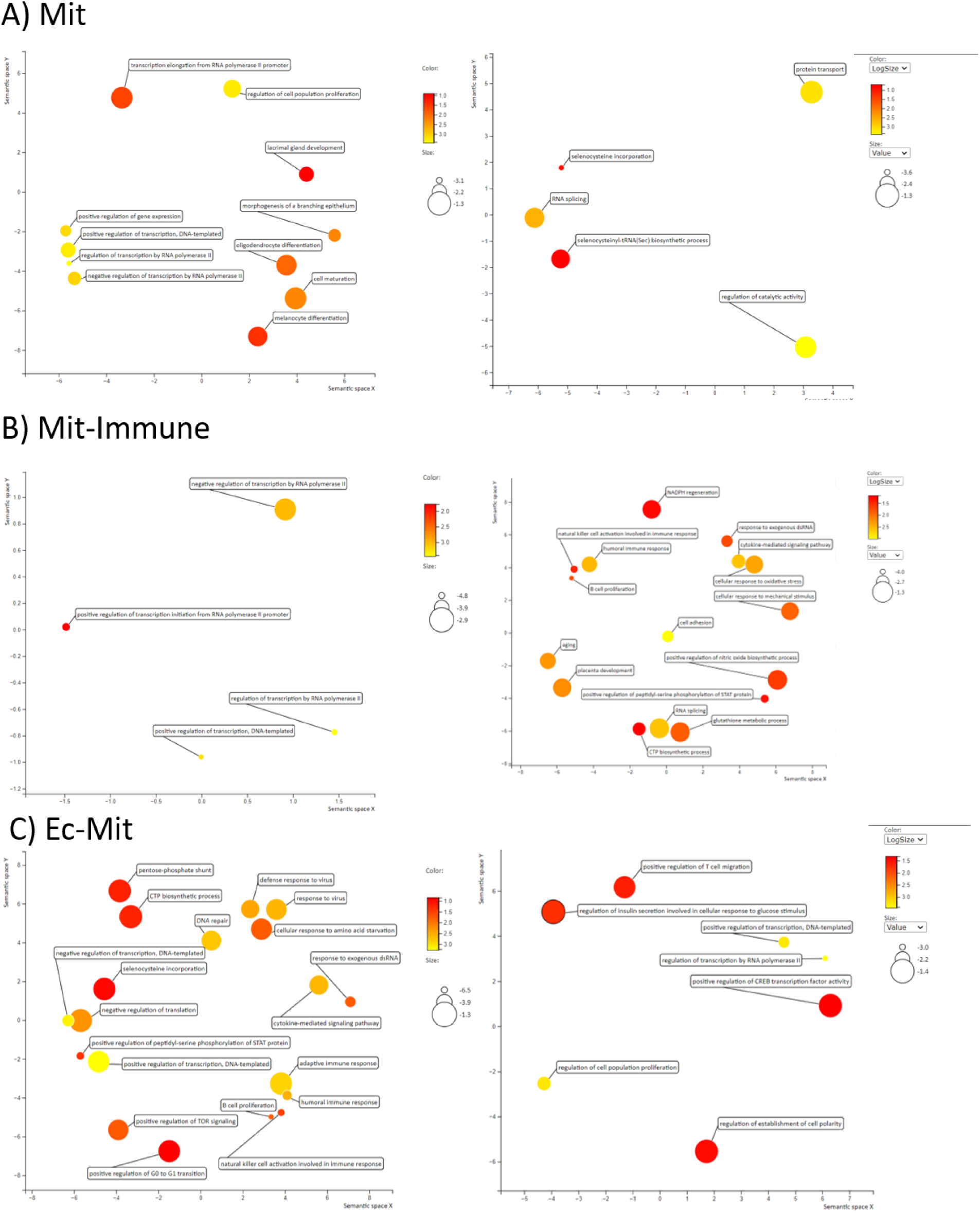
Pathway analyses (DAVID, followed by REVIGO) of genes that are predicted be targetable in MIT (a), MitImmune (b) and ECMIT (c) high tumors. Note, another interpretation of the Figure: Pathway analyses of genes that are predicted to show genetic <u>resistance</u> in MIT (a), MitImmune (b) and ECMIT (c) <u>high</u> tumors.

**Figure S4.**
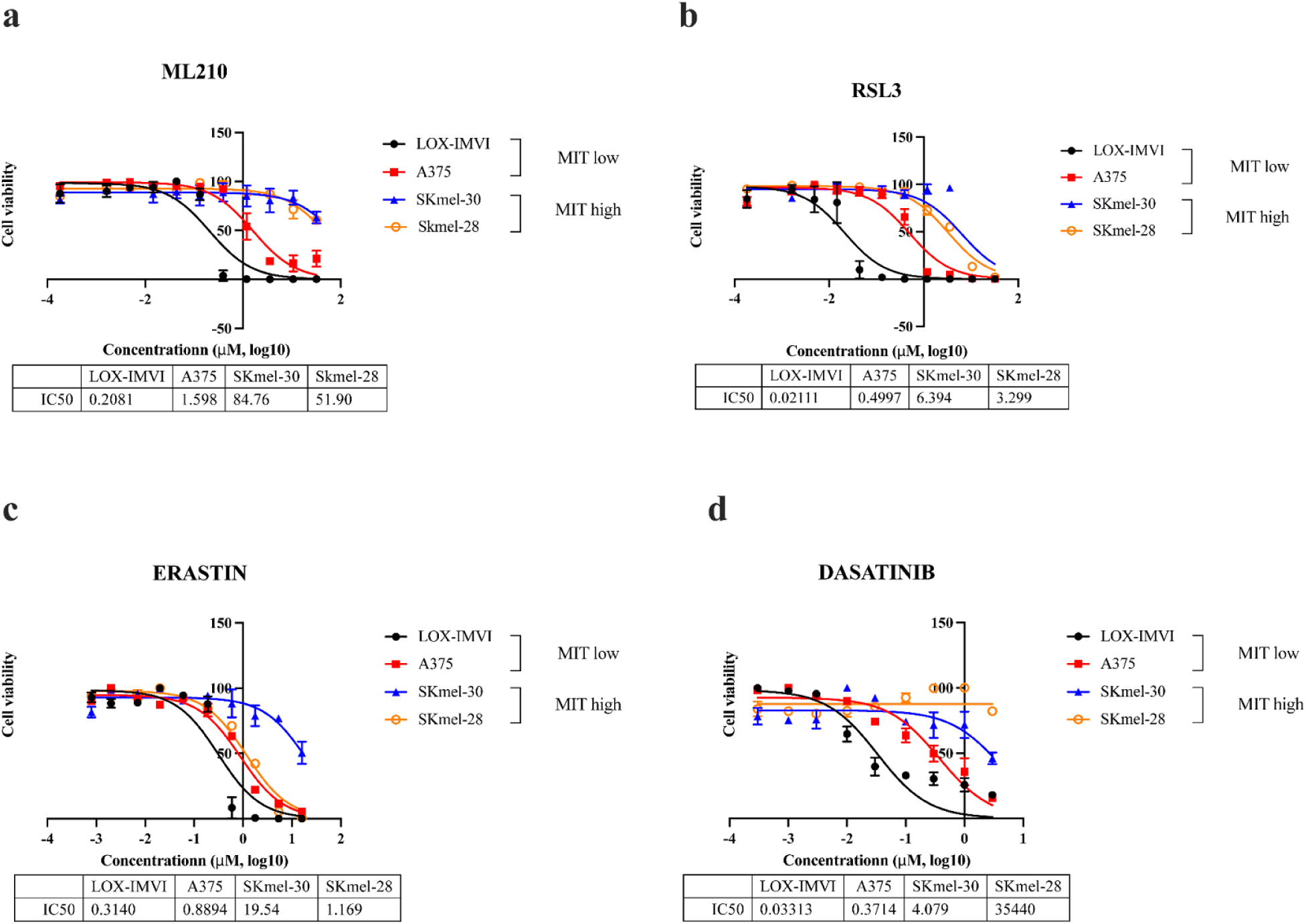
In vitro validation experiments demonstrated increased sensitivity of MIT low melanoma cells to ferroptosis inducer ML210 (a), RSL3 (b), erastin (c) and to dasatinib (d).

